# Effect of electrical hybrid-frequency waterbath stunning on the spontaneous electroencephalogram (EEG) and electrocardiogram (ECG) of broilers

**DOI:** 10.1101/2021.08.02.454822

**Authors:** L.T La Vega, D Sato, L. V Piza, E. J. X Costa

**Affiliations:** F&S Animal Origin Food Consulting, São Paulo – SP, Postal Code 04532-060; Laboratory of Applied and Computational Physics, ZAB, FZEA, University of São Paulo – Pirassununga/SP, Postal Code 13635-900 –

**Keywords:** poultry, unconsciousness, electrical stunning, animal welfare

## Abstract

Concerns about animal welfare and meat quality have encouraged research on new methods for the stunning of broilers during animal slaughter. In this study, the electroencephalogram (EEG) and electrocardiogram (ECG) of broilers were acquired during stunning using an electrical hybrid instead of a single frequency. Considering a square-wave with a current of 220 mA and a frequency of 1100 Hz (duty-cycle 50%), the hybrid-frequency waveform is obtained generating pulses at 6600 Hertz in the pulse-width phase. Sixty broilers aged 42 days were randomly sampled; thirty were used for EEG measurement and thirty for ECG measurement. For EEG measurements, the birds’ scalps were anaesthetized, and EEG electrode needles were inserted on the subcutaneous part of the occipital scalp. For ECG, the non-invasive surface electrode was used. The electrodes were connected to a digital EEG/ECG system. The results showed that the hybrid-frequency waveform system generated epileptic forms in the birds’ EEGs. Therefore, a hybrid-frequency system may present better carcass quality results, while preserving the birds’ welfare, when compared with a single frequency system use.

## INTRODUCTION

Electrical stunning is the most common method for poultry stunning prior to slaughter, but it has been questioned on animal welfare and product quality grounds. The procedure consists of passing an electric current through the birds’ brains with a magnitude sufficient to cause uncontrolled hyperpolarization of the neurons leading to unconsciousness (Berg and Raj, 2015). Animal welfare at the time of slaughter is under Regulation 1099/2009 of the European Union (European Union Council, 2009), and the regulation permits the use of different electrical stunning systems with electrical parameters that have been scientifically demonstrated. According to the regulation, water bath stunning shall be carried out for at duration of at least four seconds and with the minimum currents shown in Table 1. The World Organization for Animal Health (OIE) in its Terrestrial Animal Health Code (OIE 2019), has also recommended these electrical parameters.

**Table 1.**
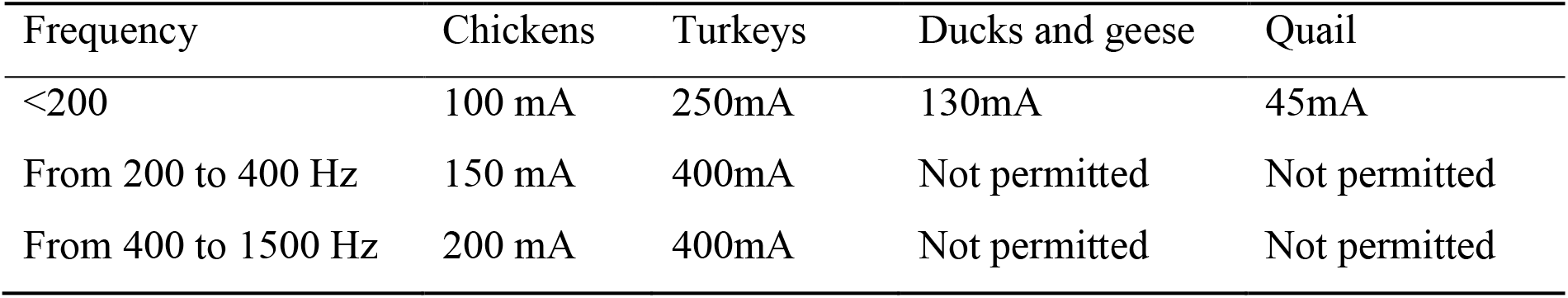
— Electrical requirements for water bath stunning equipment

| Frequency | Chickens | Turkeys | Ducks and geese | Quail |
| --- | --- | --- | --- | --- |
| <200 | 100 mA | 250mA | 130mA | 45mA |
| From 200 to 400 Hz | 150 mA | 400mA | Not permitted | Not permitted |
| From 400 to 1500 Hz | 200 mA | 400mA | Not permitted | Not permitted |

The electrical stunning equipment used to meet the aforementioned recommendations has a generator circuit that provides different waveforms, frequencies, and amplitudes of electrical currents. The equipment must have an automatic compensation circuit according to a bird’s impedance variations in the slaughter line in order to deliver a constant current to each bird. However, because the high speed of lines, this compensation cannot be performed very accurately (Berg and Raj, 2015).

Considering the guidelines of the European Union and OIE, along with the internationally designed requirements for humane slaughter, the objective of this study was to investigate electroencephalogram (EEG) and electrocardiogram (ECG) patterns of broilers stunned with hybrid electrical current frequencies, instead of single frequencies, to evaluate their impact on animal welfare. The experiment was conducted in the COPACOL slaughterhouse company, located in city of Cafelândia, in the state of Paraná, Brazil. The study was approved by the Ethics Committee on Animal Use (CEUA / USP-FZEA) under Protocol CEUA n° 4042150818. The authors declare no conflict of interest.

## MATERIAL AND METHODS

For ECG and EEG, a total of 60 broilers, ROSS line males with an average weight of 2.96 kg (std = 0.02) at 42 days-old, were used. Thirty broilers were used for EEG evaluation and 30 for ECG evaluation. Broilers were taken directly from the production line and brought to the experimental location created in the poultry sector of the COPACOL Company.

The electrical stunning equipment used, model UFX 7 (provided by Fluxo^®^ Industrial Electronics -SC / Brazil), has selection waveforms for direct or alternating electric currents (DC/AC), a variable frequency of 20-3000 Hz, duty cycle regulation from 10 to 90%, an output voltage of 10-350 VRMS (Root Mean Square Voltage), and a couple of hybrid-frequency options. The electric current amplitude was registered with a True RMS digital multimeter model U1252B (Keysight Technologies^®^ -USA), and the waveforms were monitored with a portable oscilloscope model H110-037 (HOMIS^®^ -Brazil).

### Hybrid frequency waveform system

Considering a square wave DC current with an amplitude of 220 mA ± 10 mA and a frequency of 1100 Hz (duty cycle 50%), the hybrid frequency waveform was created by the application of pulses of 6600 Hz in half of each cycle, in the pulse width time. Figure 1 depicts a graphical representation of hybrid waveform used in this study.

**Figure 1 –.**
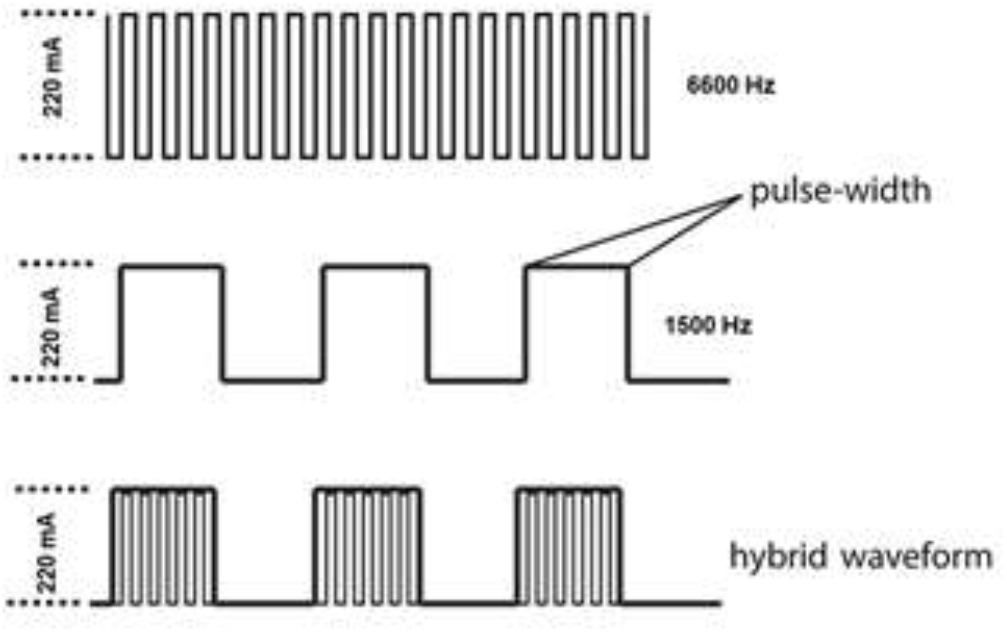
Graphical representation of hybrid frequency waveform used in the study

For ECG measurements, broilers were fitted with commercially available (2223BRQ 3M^®^ model), self-adhesive ECG electrodes, which were adhered to cleaned skin overlying the pectoralis muscles on either side of the sternum, with a ground electrode under the right leg (Figure 2).

**Figure 2 –.**
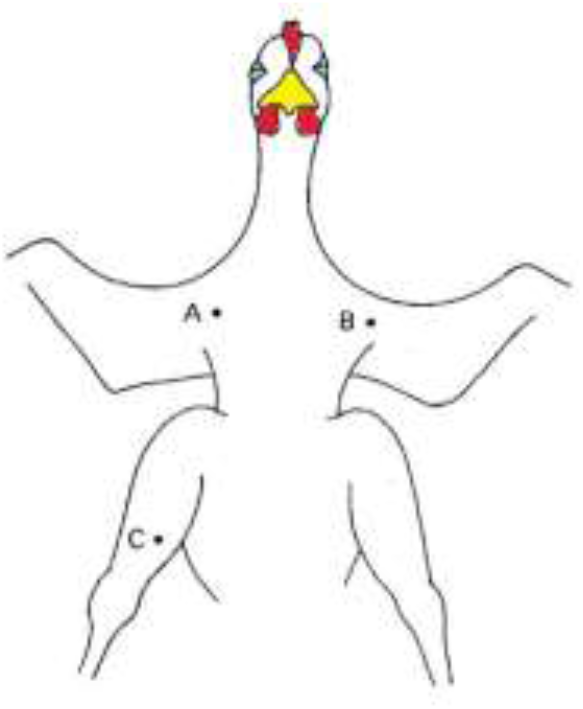
Diagram of electrode placement on the broilers

For the EEG measurements, the birds were individually implanted with needle electrodes (55% silver, 21% copper, 24% zinc of 10 mm x 1.5 mm diameter – Neurosoft^®^ mod. NS-NE-P-250/13/04) positioned under the skullcap, through the skin and skull onto the brain lobes (0.3 cm left and right of the sagittal suture and 0.5 mm toward an imaginary transverse line at the caudal margin of the eyes), with one reference electrode placed on the right or left leg. The birds were anesthetized with lidocaine (0.5 ml) applied subcutaneously on the same electrode areas, by using 31 gauge BD syringes (BD Ultra-Fine™). The electrodes were connected to the broilers with press-stud electrical connections.

To assess the EEG and ECG signals, each broiler was wrapped with a containment constructed Lycra harness, which was fastened using Velcro. The harness had a pocket to hold the wires from the electrodes and the EEG or ECG readers. The wrapped birds were placed on the slaughter line shackling and underwent the normal stunning process. The electrical water bath had the capacity for 12 birds at a time. As soon as the bird left the electrical water bath, the equipment (ECG/EEG) was automatically turned on, and the bird was taken off the line and hung on an auxiliary shackling line. The EEG/ECG signals were acquired for 60 seconds before and 60 seconds after the stunning process using wireless technology.

The EEG and ECG were collected at a sample frequency of 120 Hz and 200 Hz, respectively; the digital signal obtained was processed using fast Fourier transform (FFT), implemented using the MATLAB tool in order to obtain the frequency spectrum of different stretches associated with different events on the EEG (De Sousa Silva *et al*. 2005). Several successive artifact-free stretches were also analyzed and filtered using elliptical filters integrated into a visual tool developed at MATLAB^®^.

## RESULTS & DISCUSSION

The most important electrical parameters in electrical stunning are the current, voltage, frequency, and resistance. The electrical current, measured in amperes (A), is defined as the amount of electric charge flowing through a conductor. In electric circuits, this charge is carried by electrons and ions in an electrolyte. The electric potential difference, known as voltage, is the difference of electrical charge between two points, measured in volts (V). The electrical impedance is the measure of the difficulty of an electric current to flow through a conductor, and it is measured in ohms (Ω). In living tissues, a better term to define impedance is bio impedance (Grimnes and Martinsen, 2015). These three electrical parameters are closely related through Ohm’s law (Eq.1):

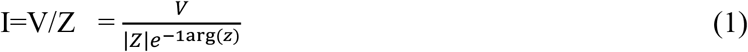

The tissue impedance can be modeled by using Equation 2 (Seoane *et al*., 2004).

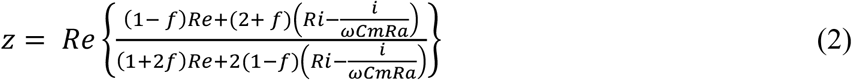

where:

Z= impedance of tissue seo

Re = resistivity of extracellular fluid (Ωcm)

Ri = resistivity of cytoplasm (Ωcm)

Cm = surface membrane capacity (Farads/cm^2^)

Ra = cell radius (cm)

F = volume factor of cell concentration

ω = angular frequency (rads/s)

i = the imaginary unity (root square of -1)

A great variation in electrical parameters has been used in electric stunning studies (Table 2). Even when the same frequency is used, for example, there are differences between voltages and waveforms, as well as amplitudes of the electric current and times of shock application. This fact may make direct result comparison challenging.

**Table 2.**
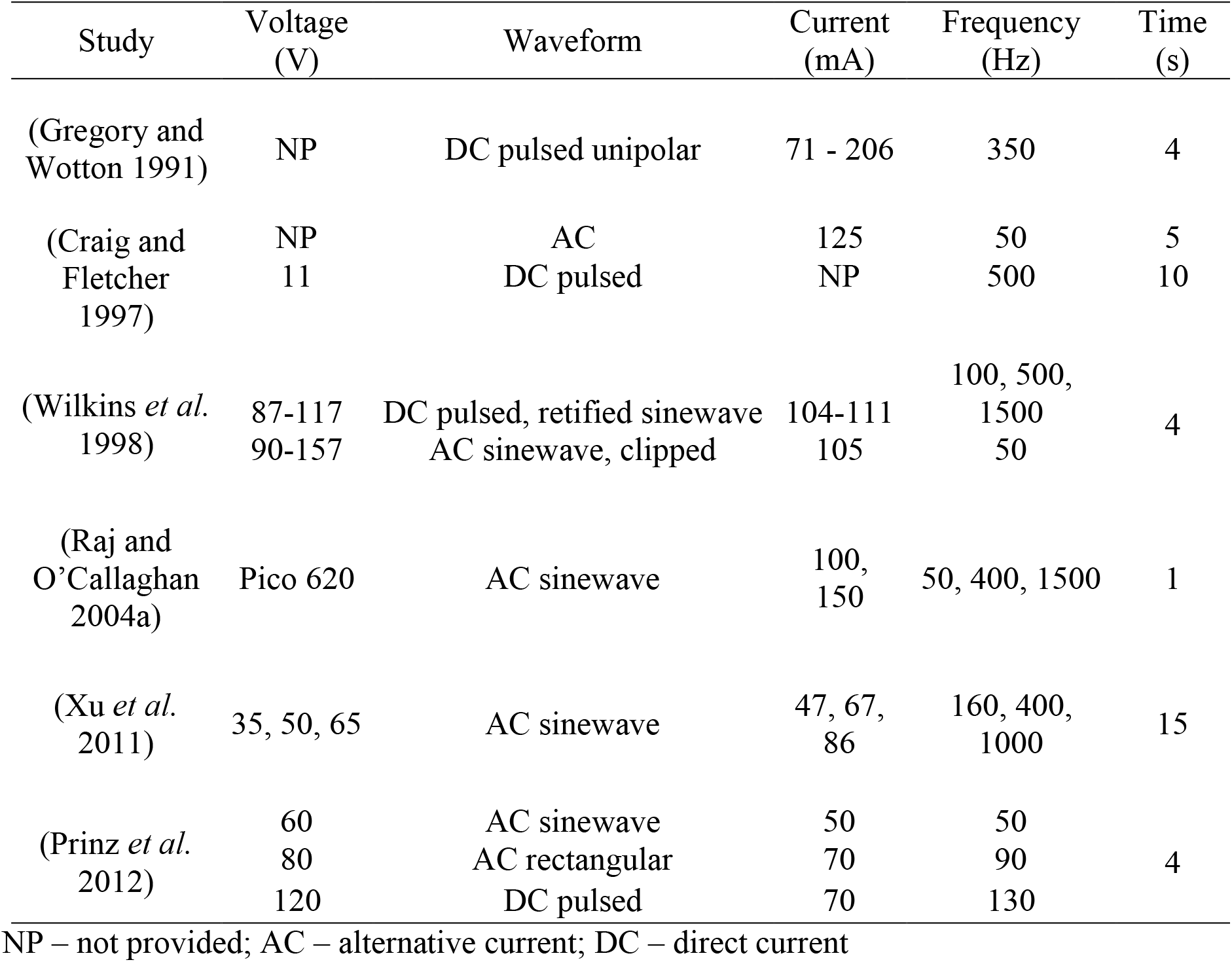
– Different waveforms, stun-time and current/voltage levels applied in electrical stunning studies

| Study | Voltage (V) | Waveform | Current (mA) | Frequency (Hz) | Time (s) |
| --- | --- | --- | --- | --- | --- |
| (Gregory and Wotton 1991) | NP | DC pulsed unipolar | 71 - 206 | 350 | 4 |
| (Craig and Fletcher 1997) | NP | AC | 125 | 50 | 5 |
|  | 11 | DC pulsed | NP | 500 | 10 |
| (Wilkins *et al.* 1998) | 87-117 | DC pulsed, rectified sinewave | 104-111 | 100, 500, 1500 | 4 |
|  | 90-157 | AC sinewave, clipped | 105 | 50 |  |
| (Raj and O'Callaghan 2004a) | Pico 620 | AC sinewave | 100, 150 | 50, 400, 1500 | 1 |
| (Xu *et al.* 2011) | 35, 50, 65 | AC sinewave | 47, 67, 86 | 160, 400, 1000 | 15 |
| (Prinz *et al.* 2012) | 60 | AC sinewave | 50 | 50 | 4 |
|  | 80 | AC rectangular | 70 | 90 |  |
|  | 120 | DC pulsed | 70 | 130 |  |

NP – not provided; AC – alternative current; DC – direct current

According to Table 2, the parameters used in this study (220 mA ± 10 mA, 1100 Hz (Duty Cycle 50%)) to carry out the experiments have been applied previously in other studies.

### EEG response in electrically stunned chickens

Digital processing of the EGG data reveal that 100% of the birds, at the exit of the electrical water bath, had an epileptic form followed by a quiescent pattern. This pattern lasted until the moment of bleeding, presenting no EEG pattern compatible with return of consciousness; i.e., after bleeding the EEG evolved to an isoelectric pattern. In addition, the data show that no bird had brain death before cutting.

Figure 3 shows representative brain electrical activity before and after the stunning process of one of the birds, randomly selected. The same phenomenon occurred in all birds sampled.

**Figure 3 -.**
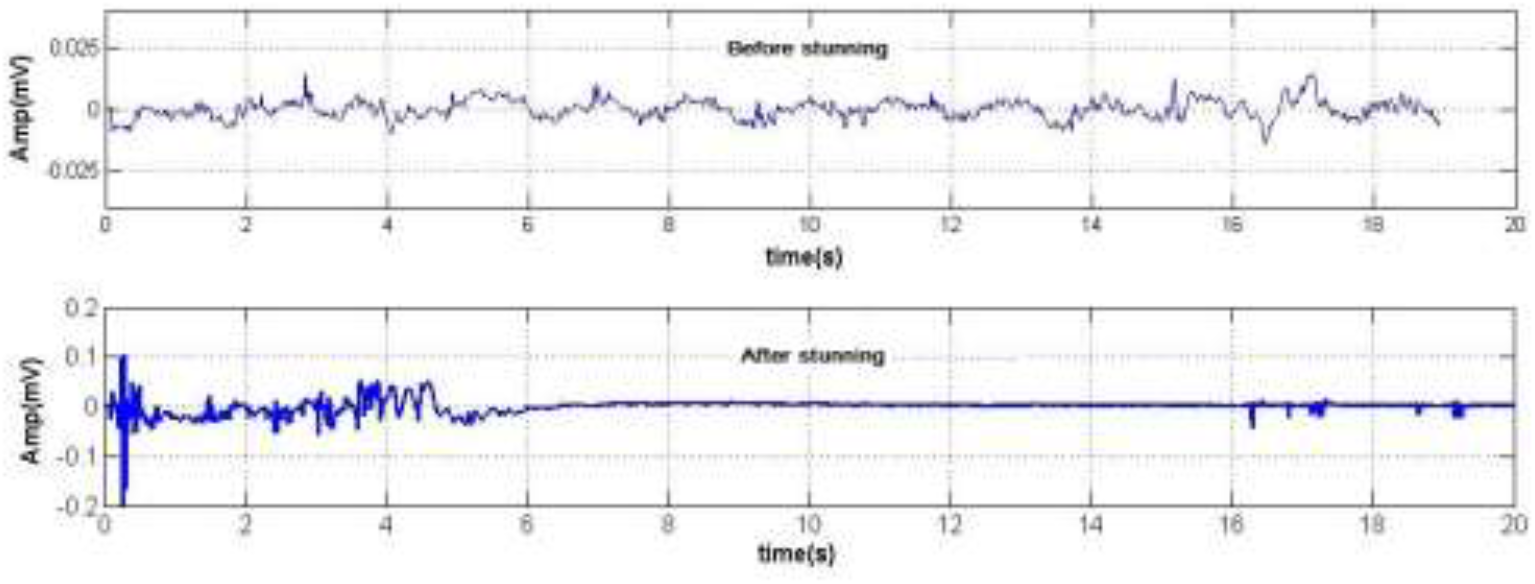
EEG pattern before and after Stunning process

For all acquired EEG signals, the signal Power Spectral Density (PSD) was calculated using the Welch Method (Manshouri *et al*., 2018). PSD displays the power distribution among the frequency components. Epileptic events have a different frequency distribution than normal brain signals, characterized mainly by an increase in frequency. Figure 4 shows the PDS calculated for EEG before and after stunning for a 95% confidence interval.

**Figure 4 –.**
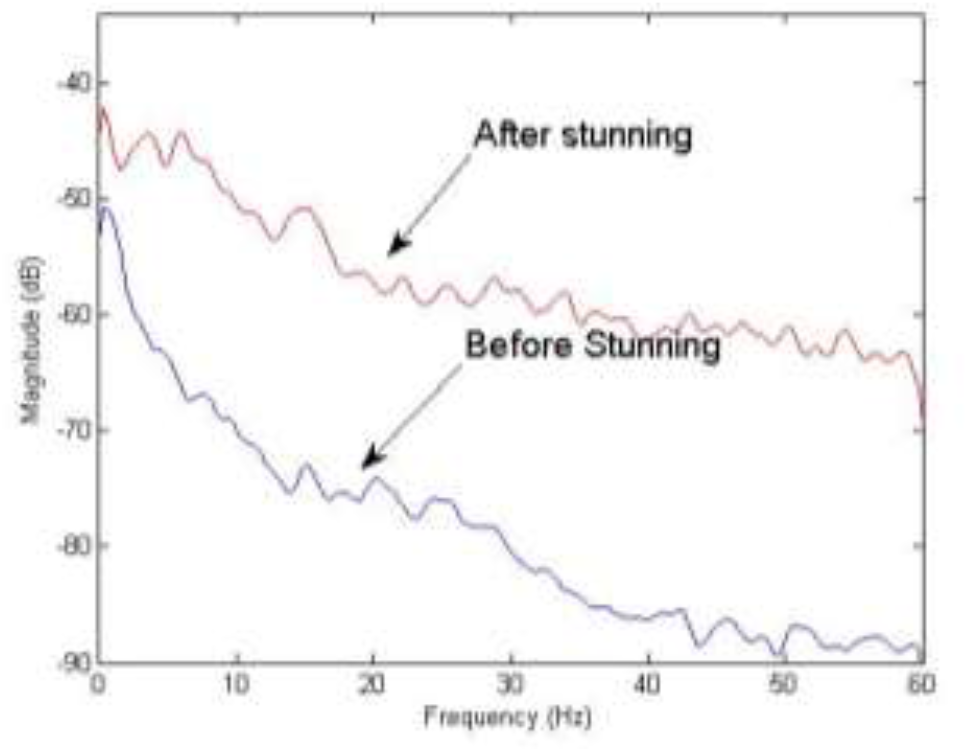
EEG Power Spectral Density (PSD) estimated using the Welch Method

In the brain, the electric current created by the shooting of the neurons is dissipated throughout the brain tissue, reaching the surface of the scalp, and can be captured and evaluated. The graphic record of the variation in the amplitude of the electrical brain activity based on time is an EEG.

The electric brain signals show features that distinguish them from other biological signs, such as those coming from the heart. According to the European Food Safety Authority (EFSA), efficient stunning methods disrupt the neurons or neurotransmitter regulatory mechanisms in the brain, causing a long-lasting depolarized neuronal state that renders animals unconscious and insensible (EFSA 2004). The EEG is the most reliable indicator of the unconsciousness and insensibility of the birds, since somatosensory reflexes and direct observations are not sufficiently reliable indicators of insensibility at high frequencies (EFSA 2013).

The use of signal processing techniques represent an important tool for identifying patterns on the EEG. The employment of digital processing methods for signals allows for more information to be obtained from the EEG signals than is obtained in its representation in the form of a time series. These processing methods include Fourier analysis, time-frequency, non-linear models, as well as the theory of complex systems and wavelets (Marchant, 2003). As seen in Figure 4, when the data are transformed, the frequencies and their magnitudes are more prominent, and the differences between before and after stunning can be better quantified.

### ECG response in electrically stunned chickens

The ECG measurement is an important physiological measurement to analyze broilers’ behavior (Blanchard *et al*., 2002). Actually, ECG had been used mainly to monitor broilers under Low Atmospheric Pressure Stunning (LAPS) methods (Martin *et al*., 2016), and few studies have been conducted on electrical water bath stunning methods. Notwithstanding, ECG can be used to monitor the broilers’ heart rates before and after electrical stunning (Barbosa *et al*., 2016).

After the ECG data were digitally processed, it was revealed that 100% of the birds were alive until bleeding. The ECG allowed the correct measuring frequencies of the birds’ heartbeats before and after the passage of the stunning electric current (Figure 5). Figure 5 shows the electrical activity of the heart, before and after the stunning process, of one of the birds randomly selected from the experimental runs.

**Figure 5 –.**
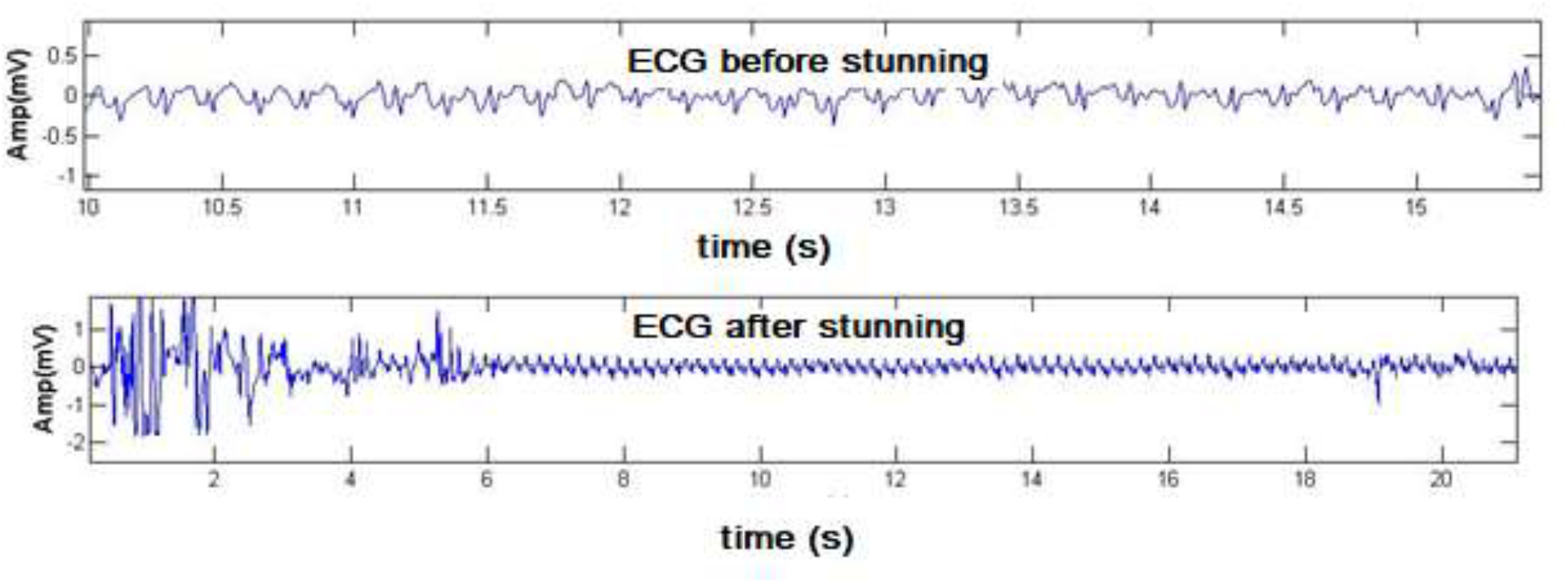
ECG signals sampled before and after the stunning process

The signals shown in Figure 5 were collected from the time the bird left the electrical water bath, i.e. just after the stunning process, and continued until after bleeding. The ECG pattern shows that, after the passage of the electric current, the heart rate, although altered in relation to frequency and amplitude, continued to represent an electrically active heart until the cut performed by the bleeder.

As suggested by other studies (Raj and O’Callaghan, 2004a; Raj *et al*,. 2006a), the depth of desensitization can be related to the pulse width time. In this case, the use of a hybrid-frequency electrical current, respecting the proportion of the single wave duty-cycle, may have contributed to obtaining increased levels of current. In living tissue, there are free ions in extracellular fluids, where the electrical current finds less resistance to flow. The lipid and protein components of living tissues offer more resistance to the passage of electrical currents, and different parameter configurations of an electrical current could change their conductivity. For example, the increase of electric current frequency has been shown to raise the conductivity of living tissues (Gabriel, 1996).

Since the use of high frequencies has the potential to improve the quality of meat, as verified by various studies (Gregory and Wotton, 1991; Wilkins *et al*., 1998; Xu *et al*., 2011; Huang *et al*., 2014), the use of a hybrid frequency wave system results in a positive effect on meat quality, without negatively affecting the animal’s welfare. However, concerning animal welfare, a very important question is whether loss of consciousness occurs instantly in electrical stunning.

The occurrence of highly synchronized 8–13 Hz activity in EEG recordings, as well as the absence of potential somatosensory evoked responses are known as reliable measurements to determine the effectiveness of stunning methods (Berg and Raj, 2015; Terlouw *et al*., 2015). In a study using different electrical single stunning frequencies, generalized epilepsy activity in an EEG of a broiler’s brain one second after electrical shock (Raj and O’Callaghan, 2004a) was observed. Despite some initial disagreement regarding the occurrence of generalized epilepsy in poultry (Gregory and Wotton, 1987; Gregory and Wotton, 1989; Raj, 2003), further research confirmed that effective electrical stunning of chickens indeed leads to epileptiform EEG activity compatible with an unconscious state (Raj and O’Callaghan, 2004b; Raj *et al*., 2006a; Raj *et al*., 2006b).

To the best of the authors’ knowledge, there is no information on the brain activity of broilers at the exact moment of electrical shock. The published studies report brain activity right after the electrical shock. The scientific knowledge about human electroconvulsive therapy without the use of anaesthetics, which includes reports of intensive pain by patients, was used in a critical comparison to the electrical stunning of broilers (Zivotofsky and Strous, 2012). Unfortunately, even in methods where animals recover after the procedure, an animal cannot express whether it had experienced pain during the electrical shock, and there is no technology available to date to answer this question.

Regarding obtaining an unconsciousness state, several studies have shown that low frequencies (50/60 Hz) are more effective, and when high frequencies are used, the amount of electrical current must be increased to maintain the effectiveness of stunning (Gregory and Wotton, 1991; Wilkins *et al*., 1998; Raj *et al*., 2006a; Prinz *et al*., 2010a; Prinz *et al*., 2010b; Prinz *et al*., 2012). Regarding meat quality, higher frequencies have shown a positive effect with a decrease of broken bones and hemorrhagic spot indices in many species (Anil and McKinstry, 1992; Turcsán *et al*., 2003; Xu *et al*., 2011; Grimsbø *et al*., 2014; Robins *et al*., 2014; Huang *et al*., 2014).

## CONCLUSIONS AND APPLICATIONS

The results of the experiments conducted at slaughterhouse of Copacol (Cafelândia -PR) with the hybrid-frequency wave current generator, using the stunning equipment UFX7 Solution^®^ support the following conclusions:

- Humanitarian slaughter: 100% of the birds that underwent electrical shock presented generalized epilepsy after being submitted to the pre-established electrical standards of the European Union and OIE. This conclusion is supported by clinical analysis and EEG data analysis.
- Religious slaughter -Halal: 100% of the birds were still alive after the electric shock according to the EEG and ECG data, as well as clinical signs. There was no impact on the killing or bloodletting process. The use of hybrid-frequency current is effective to promote humanitarian and Halal slaughter, guaranteeing epilepsy followed by a quiescence period, and keeping animals alive after desensitization. In addition, equipment with an effective combination of hybrid waves allows for adjustments, not only of the amplitude and frequency of the waves, but also to the waveform, duty cycle, and complex wave compositions. This flexibility benefits the animals, which suffer no pain, and the slaughter, which incurs greater productivity. The results of the research show that EEG and ECG data can be used to support the requirements of humanitarian and Halal slaughter, respectively.

## Acknowledgment

The authors would like to thanks to Prof.. JaciMara Baptista and Ms. Madariaga-Hopkins from UNLV-USA for English revision.

## Notes

### Competing Interest Statement

The authors have declared no competing interest.

## REFERENCES

Anil MH, McKinstry JL (1992) The effectiveness of high frequency electrical stunning in pigs. Meat Science 31, 481–491. doi:10.1016/0309-1740(92)90030-8.

Berg C, Raj M (2015) A Review of Different Stunning Methods for Poultry — Animal Welfare Aspects (Stunning Methods for Poultry). Animals 5, 1207–1219. doi:10.3390/ani5040407.

Blanchard S, Degernes L, DeWolf D, Garlich J (2002) Intermittent biotelemetric monitoring of electrocardiograms and temperature in male broilers at risk for sudden death syndrome. Poultry Science 81, 887–891. doi:10.1093/ps/81.6.887.

Craig E, Fletcher D (1997) A comparison of high current and low voltage electrical stunning systems on broiler breast rigor development and meat quality. Poultry Science 76, 1178–1181. doi:10.1093/ps/76.8.1178.

EFSA (2004) Opinion of the Scientific Panel on Animal Health and Welfare (AHAW) on a request from the Commission related to welfare aspects of the main systems of stunning and killing the main commercial species of animals. EFSA Journal 2, 45. doi:10.2903/j.efsa.2004.45.

EFSA (2013) Annual Report of the EFSA Journal 2012. EFSA Supporting Publications 10,. doi:10.2903/sp.efsa.2013.EN-418.

European Union Council (2009) ‘Regulamento (CE) N. o 1099/2009 do Conselho da União Européia de 24 de Setembro de 2009 relativo à proteção dos animais no momento da occisão.’

Gabriel SG and RWL and C (1996) The dielectric properties of biological tissues: II. Measurements in the frequency range 10 Hz to 20 GHz. Physics in Medicine and Biology 41, 2251.

Gregory NG, Wotton SB (1987) Effect of electrical stunning on the electroencephalogram in chickens. The British veterinary journal 143, 175–183. doi:10.1016/0007-1935(87)90009-1.

Gregory NG, Wotton SB (1989) Effect of electrical stunning on somatosensory evoked potentials in chickens. British Veterinary Journal 145, 159–164. doi:10.1016/0007-1935(89)90098-5.

Gregory NG, Wotton SB (1991) Effect of a 350 Hz DC stunning current on evoked responses in the chicken’s brain. Research in veterinary science 50, 250–251. doi:10.1016/0034-5288(91)90118-8.

Grimnes S, Martinsen ØG (2015) Chapter 1 - Introduction BT - Bioimpedance and Bioelectricity Basics (Third Edition). pp. 1–7. (Academic Press: Oxford) doi:http://dx.doi.org/10.1016/B978-0-12-411470-8.00001-5.

Grimsbø E, Nortvedt R, Hammer E, Roth B (2014) Preventing injuries and recovery for electrically stunned Atlantic salmon (Salmo salar) using high frequency spectrum combined with a thermal shock. Aquaculture 434, 277–281. doi:10.1016/j.aquaculture.2014.07.018.

Huang JC, Huang M, Yang J, Wang P, Xu XL, Zhou GH (2014) The effects of electrical stunning methods on broiler meat quality: Effect on stress, glycolysis, water distribution, and myofibrillar ultrastructures. Poultry Science 93, 2087– 2095. doi:10.3382/ps.2013-03248.

Manshouri N, Maleki M, Kayikcioglu T (2018) Power spectrum analysis of EEG for watching 2D &amp; 3D videos and resting state. In ‘2018 26th Signal Process. Commun. Appl. Conf.’, 1–4. (IEEE) doi:10.1109/SIU.2018.8404394.

Marchant BP (2003) Time–frequency Analysis for Biosystems Engineering. Biosystems Engineering 85, 261–281. doi:10.1016/S1537-5110(03)00063-1.

Martin JE, Christensen K, Vizzier-Thaxton Y, McKeegan DEF (2016) Effects of light on responses to low atmospheric pressure stunning in broilers. British Poultry Science 1–16. doi:10.1080/00071668.2016.1201200.

OIE (2019) ‘Terrestrial Animal Health Code 2019. Volume 1 : General provisions.’ (© OIE (World Organisation for Animal Health)) https://www.oie.int/standard-setting/terrestrial-code/.

Prinz S, Van Oijen G, Ehinger F, Bessei W, Coenen a (2010) Effects of waterbath stunning on the electroencephalograms and physical reflexes of broilers using a pulsed direct current. Poultry Science 89, 1275–1284. doi:10.3382/ps.2009-00136.

Prinz S, Van Oijen G, Ehinger F, Bessei W, Coenen a. (2012) Electrical waterbath stunning: Influence of different waveform and voltage settings on the induction of unconsciousness and death in male and female broiler chickens. Poultry Science 91, 998–1008. doi:10.3382/ps.2009-00137.

Prinz S, Van Oijen G, Ehinger F, Coenen A, Bessei W (2010) Electroencephalograms and physical reflexes of broilers after electrical waterbath stunning using an alternating current. Poultry Science 89, 1265–1274. doi:10.3382/ps.2009-00135.

Raj a BM (2003) A critical appraisal of electrical stunning in chickens. World’s Poultry Science Journal 59, 89–98. doi:10.1079/WPS20030005.

Raj ABM, O’Callaghan M (2004a) Effect of amount and frequency of head-only stunning currents on the electroencephalogram and somatosensory evoked potentials in broilers. Animal Welfare 13, 159–170.

Raj a BM, O’Callaghan M (2004b) Effects of electrical water bath stunning current frequencies on the spontaneous electroencephalogram and somatosensory evoked potentials in hens. British poultry science 45, 230–236. doi:10.1080/00071660410001715830.

Raj ABM, O’Callaghan M, Knowles TG (2006a) The effects of amount and frequency of alternating current used in water bath stunning and of slaughter methods on electroencephalograms in broilers. Animal Welfare 15, 19–24.

Raj ABM, O’Callaghan M, Knowles TG (2006b) The effects of pulse width of a direct current used in water bath stunning and of slaughter methods on spontaneous electroencephalograms in broilers. Animal Welfare 15, 25–30.

Robins a., Pleiter H, Latter M, Phillips CJC (2014) The efficacy of pulsed ultrahigh current for the stunning of cattle prior to slaughter. Meat Science 96, 1201–1209. doi:10.1016/j.meatsci.2013.10.030.

De Sousa Silva AC, Céspedes Arce AI, Souto S, Xavier Costa EJ (2005) A wireless floating base sensor network for physiological responses of livestock. Computers and Electronics in Agriculture 49,. doi:10.1016/j.compag.2005.05.004.

Terlouw C, Bourguet C, Deiss V (2015) Consciousness, unconsciousness and death in the context of slaughter. Part II. Evaluation methods. Meat Science. doi:10.1016/j.meatsci.2016.03.010.

Turcsán Z, Varga L, Szigeti J, Turcsán J, Csurák I, Szalai M (2003) Effects of electrical stunning frequency and voltage combinations on the presence of engorged blood vessels in goose liver. Poultry science 82, 1816–1819.

Wilkins LJ, Gregory NG, Wotton SB, Parkman ID (1998) Effectiveness of electrical stunning applied using a variety of waveform-frequency combinations and consequences for carcase quality in broiler chickens. British poultry science 39, 511–518. doi:10.1080/00071669888692.

Xu L, Zhang L, Yue HY, Wu SG, Zhang HJ, Ji F, Qi GH (2011) Effect of electrical stunning current and frequency on meat quality, plasma parameters, and glycolytic potential in broilers. Poultry science 90, 1823–1830. doi:10.3382/ps.2010-01249.

Zivotofsky AZ, Strous RD (2012) A perspective on the electrical stunning of animals: Are there lessons to be learned from human electro-convulsive therapy (ECT)? Meat Science 90, 956–961. doi:10.1016/j.meatsci.2011.11.039.

